# Evolution of mutation rates in digital genomes: the roles of genetic drift, mutational supply, and genome size

**DOI:** 10.64898/2026.07.03.736272

**Authors:** Quentin Fernandez de Grado, Antoine Frénoy

## Abstract

Mutation is the ultimate mechanism that produces genetic novelty, and thus a central ingredient of evolution. Mutation rates are therefore thought to be tuned by natural selection, for example to optimize a delicate balance between the generation of adaptive diversity and the accumulation of deleterious mutations.

As this selection occurs over very long time scales, models and simulations have been powerful tools to understand how mutation rate evolves and which factors influence it. Most simulation methods are nevertheless limited by the over-simplicity of the genotype-to-phenotype map they feature, especially regarding the encoding of mutation rate.

We modified Aevol, an evolutionary simulator inspired by bacterial genomics with a realistic genome structure and a complex genotype-to-phenotype layer, to allow organisms to evolve genes coding for higher replication fidelity. This setup permits several degrees of realism absent in other models: mutation-rate modifier genes themselves experience a realistic distribution of effects of mutations and diminishing-returns epistasis, similarly to fitness modifiers. Moreover, a lower mutation rate comes with the trade-off of a larger genome to encode the genes improving replication fidelity.

We use this setup to test hypotheses regarding the evolution of prokaryotic mutation rate, and its link with genome size and genetic drift. We found that evolution systematically increases replication fidelity, even when this results in lower fitness. We highlight two factors which limit the mutation rate decrease: genetic drift and the supply of gain-of-fidelity mutations.

**Significance Statement:** Mutation rate is a central parameter governing the evolution of living systems, but it is also itself the product of evolution, as it is determined by enzymatic processes which are subject to hereditary variations and natural selection. Several hypotheses exist to explain how the mutation rate evolves, and which factors govern mutation rate variation between and within species. We propose a “digital genomics” simulation model which permits testing and refining some of these hypotheses, in a setup capturing key constraints such as a realistic supply of mutations and selection pressure for genome space. We found that selection almost always decreases the mutation rate. We highlight the role of two factors in determining the amount of mutation rate reduction, genetic drift and the supply of gain-of-fidelity mutations, as well as a strong relationship with genome size.

## Introduction

Mutation is the primary mechanism of evolution that produces genetic novelty and grants living organisms the capacity to evolve. However, because such novelty is mostly deleterious and rarely beneficial, living organisms have developed biological systems that increase the fidelity of DNA replication and reduce the rate of mutations. Since these fidelity systems are themselves encoded on the genome, the mutation rate of living organisms is not set in stone and can evolve. Indeed, genes contributing to replication fidelity are themselves subject to mutations (Treffers 1954) and selection (Chao and Cox 1983).

Several non-exclusive scenarios have been proposed to explain how the mutation rate evolves. The simpler scenario states that, because most mutations are deleterious, selection always decreases mutation rate (Liberman and Feldman 1986; Williams 1996). This decrease can reach a biochemical limit, or alternatively can be limited by the costs associated with fidelity systems, such as energy consumption, time delay, and genomic space used to encode them (Kimura 1967; Kirkwood et al. 1986; Kondrashov 1988; Kondrashov 1995; Furió et al. 2005). A complementary scenario, named drift-barrier theory, proposes that the main limit in mutation rate reduction is genetic drift: below a certain mutation rate threshold value determined by effective population size, the effect of an allele further decreasing mutation rate would be too small to be favored by selection (Lynch 2011). An alternative scenario considers the importance of adaptive mutations and postulates that selection for evolvability can also favor alleles which increase mutation rate (Chao and Cox 1983; Taddei et al. 1997). Therefore long-term selection would push the mutation rate toward an intermediate value that balances the constraints of minimizing the mutational load with the need for adaptive mutations to cope with environmental changes (Kimura 1967, Levins 1967, Leigh 1970, Eshel 1973, Gillespie 1981, Ishii et al. 1989, Bedau and Packard 2003, Clune et al. 2008, Good and Desai 2016).

As this indirect selection for mutation rate happens over very long time scales, models and simulations have been powerful tools to understand how mutation rate evolves and which factors influence this evolution. However, the complex, second-order pressures for robustness and for evolvability which act on mutation rate evolution are not easy to incorporate in models, as they depend on parameters that are hard to estimate, such as the distribution of fitness effects of mutations (Bataillon and Bailey 2014), and the way several mutations combine (epistasis, Martin et al. 2007).

These models greatly vary in nature (analytical, simulations) and in levels of complexity, but share the same general conception: individuals are characterized by their fitness (which can be determined by one or several loci for models featuring an explicit genetic representation), and by their mutation rate, classically encoded by a modifier locus which does not directly affect the fitness but the mutation rate at other loci, and is under second-order selection (Kimura 1967; Levins 1967; Raynes et al. 2018). The genotype-to-phenotype-to-fitness map can be more or less complex, ranging from individuals represented as a single floating point fitness value with fixed effects of mutations (Taddei et al. 1997) to digital genomes interpreted by rules leading to complex fitness landscapes with emergent fitness effects of mutations (Clune et al. 2008). Most models fall on a middle-ground between these two extremes, for example without an explicit genomic representation but with realistic and finely tunable fitness effects of mutations (Good and Desai 2016).

Small variations of this general pattern have been proposed, for example to consider a direct cost of high replication fidelity (Ishii et al. 1989; André and Godelle 2006), or to explicitly represent resources and simulate agents movement and feeding to determine reproduction (Bedau and Packard 2003).

We consider several limitations shared by most of these models, which motivate the current study in which a simulation system able to bypass these limitations is considered:

### Mutation rate encoding

While some of the existing models permit a complex genotype-to-phenotype-to-fitness map with a realistic distribution of fitness effects of mutations, this map is not applied to mutation rate themselves, which are generally simply encoded as a floating point value which can evolve upwards and downwards (often but not necessarily with equal probabilities). In organic life, genes increasing replication fidelity are themselves facing similar distributions of effects of mutations than genes directly contributing to fitness, namely most mutations are loss-of-functions, that is to say increase mutation rate, and mutations that decrease mutation rate should be rare. Additionally, for mutation rate as for other traits, we can expect epistasis (non-additivity of the effects of several mutations), and diminishing returns (organisms that have evolved a low mutation rate have less accessible mutations which further decrease mutation rate).

### Genome size and cost of genomic information

While some models can artificially associate a cost to lower mutation rates (for example by representing the associated increase in replication time, or the metabolic cost associated with the enzymatic repair), they can not consider the intrinsic cost of carrying more genetic information. In organic life, genome sizes are thought to be tightly controlled (Drake 1991; Drake et al. 1998; Knibbe et al. 2007), for example because larger genomes are subject to more mutations (in particular more rearrangements) which are likely to be deleterious. Using genomic space to encode fidelity genes thus raises an interesting tradeoff: since space is constrained, it comes with the indirect cost of decreasing the genomic space available for other traits which directly contribute to fitness; but lowering mutation rate alleviate the cost associated with larger genomes.

In this study, we choose to use Aevol, an evolutionary simulator, which has already been successfully used in the past to study the selection pressures acting on the structure of microbial genomes (Knibbe et al. 2007, Banse et al. 2024, and Luiselli et al. 2024). Its strengths dwell in its explicit genomic representation and complex genotype-to-phenotype map, with emergent fitness effects of mutations rather than a predefined distribution.

We modified Aevol to incorporate a self-evolved mutation rate. We did so by allowing organisms to encode fidelity genes in their genome, in addition to the metabolic genes which directly contribute to the phenotype and fitness in the original version of Aevol. The same encoding and phenotype computation rules are used for fidelity genes than for metabolic genes. The only difference is that a part of the space of traits is reallocated to fidelity, and the genomic content thus defines both the fitness and the mutation rate of an individual.

Compared to the other models discussed above, this means that there is a variable, evolvable number of fidelity genes which together contribute non-additively to the reduction of mutation rate. A given mutation rate can be attained through various combination of fidelity genes, and these fidelity genes are subject to the same emergent distribution of effects than metabolic genes.

We use this model to study evolution of mutation rate in this more realistic setup, characterized by a cost for genomic space and a complex genotype-to-phenotype map for both the fitness and the mutation rate.

## Results

Aevol features digital organisms characterized by a genome composed of a string of base-pairs, from which coding segments are determined following motif-based rules inspired from bacterial genomics. These coding segments are compiled into a phenotype, represented as an abstract list of traits associated with a performance level. In our version, these traits fall into two categories, metabolism or fidelity. Metabolism is under direct selection, as it determines the fitness of the organism, which is used to calculate the probability of this organism to be present at the next generation, whereas fidelity is under indirect selection, because it determines the mutation rate, which is the probability of a base-pair to mutate at each reproduction event.

We evolved populations of 1024 individuals for 4 million generations, with the ability to reduce their mutation rate by encoding fidelity genes. The mutation rate of naive organisms which do not encode fidelity genes is named *Basal Mutation Rate* (BMR). Fidelity genes permit to decrease mutation rate below this value (see Material and Methods for details). We tested seven different values for BMR ranging from 10^-4^ to 10^-7^ , with ten replicate simulations for each value. In parallel, as a control, we performed the same simulations, except that individuals did not have the ability to reduce their mutation rate. Fidelity genes could still evolve, but were not assigned any effect on mutation rate, which remained constantly equal to the BMR value.

### Are populations decreasing their mutation rate when allowed to do so?

In all conditions except when the basal mutation rate is already extremely low (BMR = 10^-7^ ), the populations evolved a reduced mutation rate (Figure 1a), by encoding fidelity genes (Figure 1b).

**Fig. 1:**
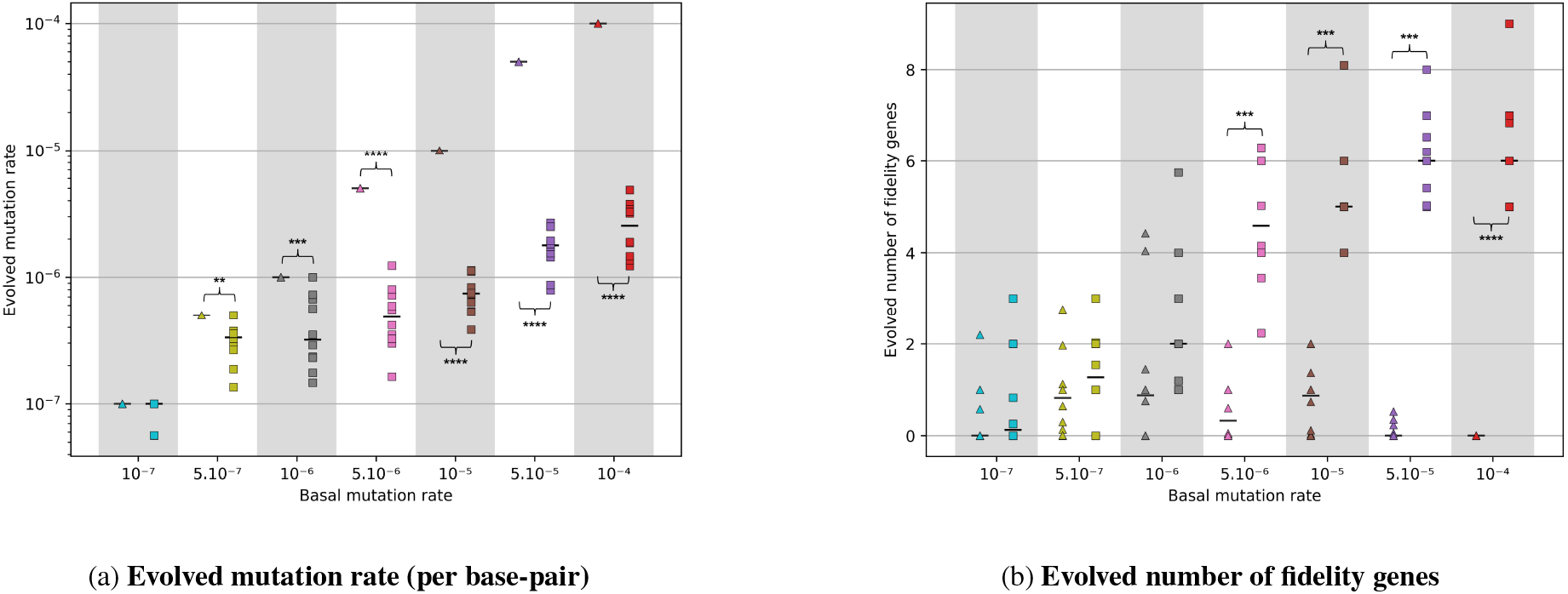
Populations evolved a lowered mutation rate by encoding fidelity genes. Populations either have the ability to reduce their mutation rate (□) or do not have it (△). Seven basal mutation rate (BMR) values are tested ranging from 10^-4^ to 10^-7^. For each BMR and each treatment, 10 replicates simulations are performed. Each simulation lasted 4 million generations. Each data point corresponds to the mean for a replicate simulation during the last 100,000 generations (see data display in Model and Methods for more details), and the horizontal bars represent the median among replicate simulations. A Mann–Whitney U test is performed and only significant p-values are reported (⋆ = p-value ≤ 0.05, ⋆⋆ = p-value ≤ 0.01, ⋆⋆⋆ = p-value ≤ 0.001, and ⋆⋆⋆⋆ = p-value ≤ 0.0001).

We can make two observations on the mutation rate; first, the evolved mutation rates do not tend toward the same value across the BMR conditions, and second, the mutation rate decrease is bigger when the BMR value increases.

The mutation rate decrease is related to the number of fidelity genes encoded, even through each fidelity gene can contribute at different degree to the decrease. Some control populations also encoded a few fidelity genes, which have no effect on the phenotype and are only subject to mutations and genetic drift. Comparing the number of fidelity genes evolved in the control populations with fixed mutation rate and in the “treatment” populations which could decrease their mutation rate permits to infer whether the mutation rate decrease is selected or can be solely explained by mutations and genetic drift.

For medium and high BMR values (10^-4^ , 5.10^-5^ , 10^-5^ and 5.10^-6^ ), we observe a significant difference between the number of fidelity genes encoded in the control populations and in the “treatment” populations (Figure 1b). When the BMR values are lower (10^-6^ , 5.10^-7^ and 10^-7^ ), the number of fidelity genes encoded are not significantly different, although mutation rates differ significantly for BMR values 10^-6^ and 5.10^-7^.

### Why do populations decrease their mutation rate, and what is the link with genome size?

A higher mutation rate increases both the supply of beneficial and deleterious mutations. Our finding suggest that in our setup, selection pushing down mutation rate to decrease mutational load predominates over selection for increasing mutation rate to increase evolvability, in line with the reduction principle (Liberman and Feldman 1986) proposed in the literature for sufficiently stable environments.

An inverse correlation between mutation rate and genome size has been empirically observed in prokaryotes (Drake 1991; Drake et al. 1998; Bourguignon et al. 2020), and a part of the literature suggests that this selection for decrease mutational load explains this inverse correlation: a higher mutation rate would select for a decrease in genome size to decrease the total number of mutations that individuals undergo (Knibbe et al. 2007; Luiselli et al. 2024).

Our observations are in agreement with this line of thought: (1) We found that in the control, when mutation rate is constant and equal to the BMR, individuals evolved smaller genomes when the BMR value increases (Figure 2a, triangles); and (2) We found that “treatment” populations which reduced their mutation rate evolved larger genomes (Figure 2a, squares) than the control populations.

**Fig. 2:**
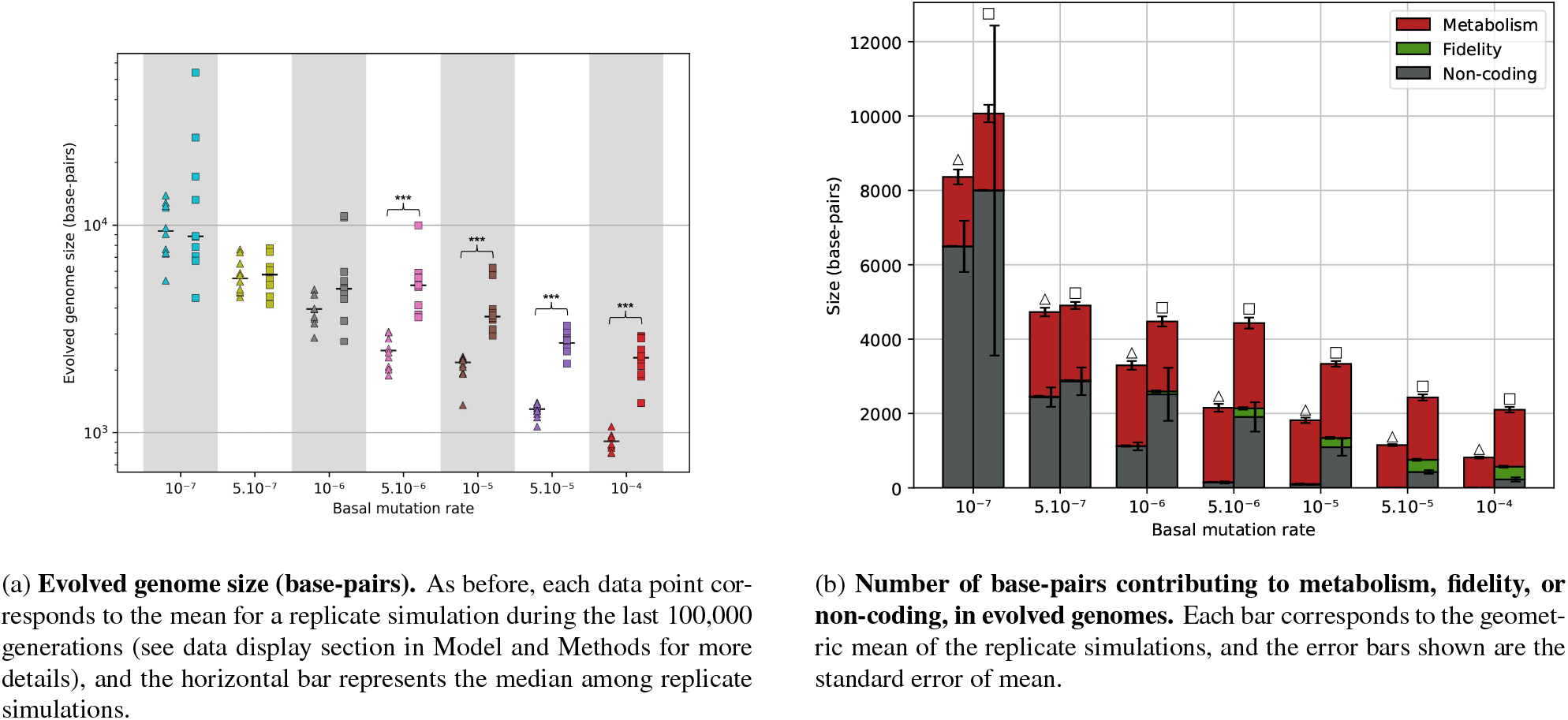
Evolved genome size, and number of base-pairs contributing to metabolism, fidelity, and non-coding. The populations are the one from Fig 1, which either have the ability to reduce their mutation rate (□) or do not have it (△).

Interestingly, Drake 1991 did not only suggest an inverse correlation between genome size and per base-per mutation rate, but also that the product of both, which is the total number of mutation per genome division, is almost constant. We make an almost similar observation of near-constant per genome mutation rate for individuals which could evolve replication fidelity, much more than for populations which could not (Figure S2).

We also looked at the amount of non-coding DNA, as it was shown in Aevol that a higher mutation rate is not only correlated with smaller genomes, but also more compact genomes with less non-coding DNA (Knibbe et al. 2007; Luiselli et al. 2024). For control populations with a fixed mutation rate, the amount of non-coding base-pairs decreases to almost zero when the BMR value increases (Figure 2b, triangles, grey), which reflects a strong selection pressure to reduce genome size. However, “treatment” populations which reduced their mutation rate accumulate significantly more non-coding DNA (Figure 2b, squares, grey), which shows that the pressure on genome size is weaker.

When the BMR value is high (10^-4^ and 5.10^-5^ ), the pressure on the genome size is so strong that it constrains the amount of metabolism that can be encoded, and impact the evolved fitness. Such observation is very similar to the error threshold (Eigen 1971), which is the critical mutation rate before newly encoded genetic elements are destroyed more frequently than selection can reproduce them. Here, control populations are very close from this error threshold, and mutations which increase genome size will likely be discarded by natural selection even if they increase fitness by encoding new metabolic genes. We observe that “treatment” populations which decreased their mutation rate are able to encode more metabolic genes (Figure S1a) and allocate more base-pairs to the metabolism (Figure S1b) without crossing this error threshold, and therefore evolve a higher fitness (Figure 3) than control populations with a fixed mutation rate.

**Fig. 3:**
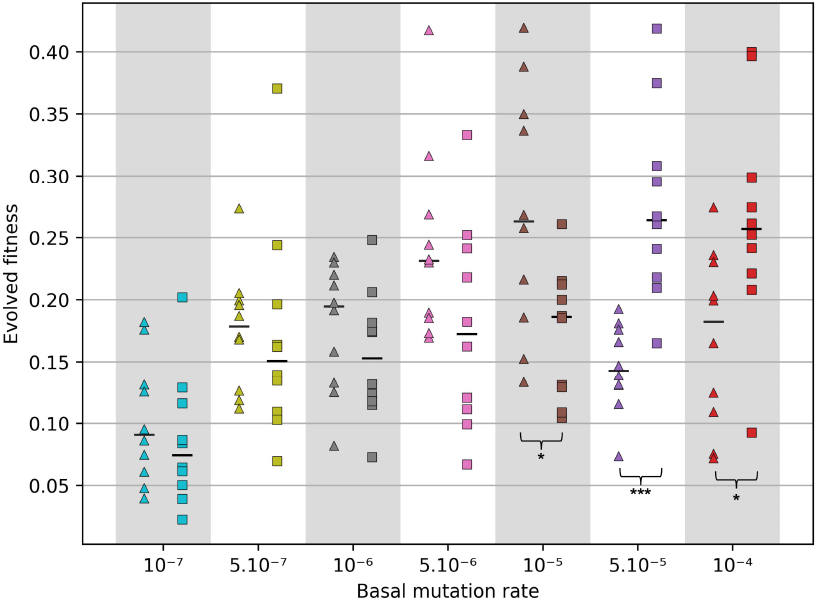
Evolved fitness in these populations from Fig 1 and 2, which either have the ability to reduce their mutation rate (□) or do not have it (△). As before, each data point corresponds to the mean for a replicate simulation during the last 100,000 generations, and the horizontal bar represents the median among replicate simulations.

When the BMR value is lower (10^-5^ , 5.10^-6^ and 10^-6^ ), the pressure on the genome is weaker and there are less constraints on the amount of metabolism that can be encoded. We can see that the number of metabolic genes and the amount of base-pairs allocated to the metabolism are similar between control populations with a fixed mutation and “treatment” populations which decreased their mutation rate (Figures S1a and S1b). However, to our surprise, in these conditions control populations with a fixed mutation rate evolved a higher fitness than treatment populations (Figure 3). A plausible explanation is that, treatment populations which decreased their mutation rate have a lower mutational supply than control populations with a fixed mutation rate, so they produce less beneficial mutations and have a lower evolvability. Indeed, as shown in Figure S2, the genomic mutation rate (defined as mutation rate per base pair per generation multiplied by genome size) is higher for control populations with a fixed mutation rate compared to “treatment” populations which decreased their mutation rate.

### Does selection systematically pushes down the mutation rate toward a sub-optimal value?

We-6described above that for BMR values 10^-5^, 5.10^-6^ and 10^-6^ , the evolved fitness is higher for control populations with a fixed mutation rate than “treatment” populations which reduced their mutation rate; thus, selection pushes the mutation rate toward a sub-opt-i4mal value for adaptation. For higher BMR values (10^-4^ and 5.10^-5^ ), where we reported that the mutation rate decrease resulted in an increased fitness compared to the control, the spontaneously evolved mutation rate may still be suboptimal, in the sense that an intermediate value between the BMR and the evolved value may have resulted in a higher fitness.

To test this, we reproduced the previous simulations for BMR values 10^-4^ and 5.10^-5^ , but we added a lower bound to the mutation rate, which is set to one order of magnitude below the BMR value, and is still higher than the evolved values when mutation rate can freely evolve.

When the mutation rate is lower-bounded, the evolved mutation rate reaches it and stays there (Figure S3a), and the evolved fitness is (slightly) higher compared to when the mutation rate can freely evolve and go below the lower bound (Figure S3b). This shows that spontaneously evolved mutation rates were also sub-optimal with these high BMR.

This is in line with the findings from Clune et al. 2008 that natural selection pushes the mutation rate toward a sub-optimal value on rugged fitness landscapes. Their proposed explanation is that on this type of landscapes, the short-term benefit of producing less deleterious mutations is stronger than the long term benefit of producing more beneficial mutations. We can think about another possible explanation in our setup, which is that fidelity genes decrease their own probability of being removed by mutations, and would therefore tend to accumulate on the genome even when detrimental to long-term adaptation, akin to selfish genetic elements.

To conclude, in all BMR conditions, selection pushes the mutation rate to a lower value than the optimal one which would confer a maximal fitness at the end of the simulation.

### What evolutionary forces prevent the mutation rate from decreasing further?

So far, we observed that selection always pushes the mutation rate downward, as stated by the reduction principle (Liberman and Feldman 1986), and this even when fidelity systems are associated with an important cost for genomic space (for high BMR values, each additional base-pair increasing the opportunity for deleterious mutations). However, we observed that neither the final evolved mutation rates nor the amount of reduction is constant between BMR conditions. We thus asked which factors determine the evolved mutation rate or the amount of mutation rate reduction. A recent hypothesis which received support from theoretical models and molecular data is the drift-barrier theory, which states that the ability of natural selection to decrease mutation rate is mostly limited by genetic drift: beyond a certain fidelity threshold, a mutation further decreasing mutation rate would have a too small effect to be selected. This threshold would depend on effective population size, which directly determines the minimal effect that a mutation should have to be visible to natural selection despite drift. This theory predicts a reverse correlation between effective population size and mutation rate, which is indeed what is observed for all domains of life (Lynch et al. 2016).

To test whether genetic drift is the limiting factor preventing further mutation rate decrease in our setup, we performed simulations where we manipulate the relative importance of genetic drift and natural selection by modifying the census population size.

In addition to the previously mentioned simulations with populations of 1024 individuals, we tested two other population sizes, 256 and 5041 individuals; and four BMR values (10^-4^ , 10^-5^ , 10^-6^ and 10^-7^ ). We performed 10 replicate simulations for each BMR value and population size combination.

The evolved mutation rate is lower in larger populations (Figure 4a), as predicted by the drift-barrier hypothesis, but the amount of reduction is lower than predicted by this theory.

**Fig. 4:**
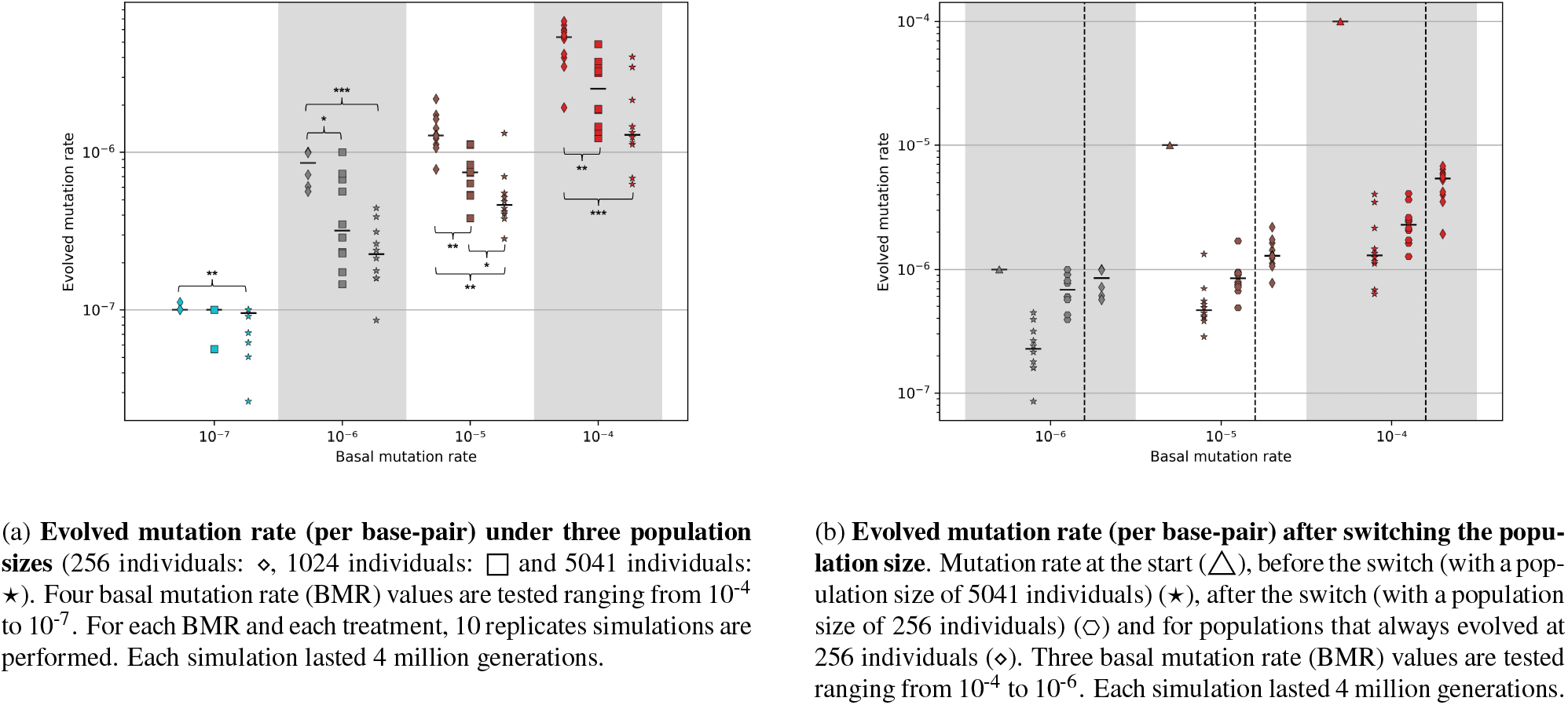
Effect of manipulating population size on mutation rate evolution. Each data point corresponds to the mean for a replicate simulation during the last 100,000 generations (see data display section in Model and Methods for more details), and the horizontal bar represents the median among replicate simulations.

However, population size does not only affect the strength of genetic drift. An alternative hypothesis could be that larger population benefit from a higher total mutational supply, including a higher supply of gain-of-fidelity alleles, providing more opportunities to improve replication fidelity. This alternative hypothesis can not be considered in population genetics models nor in simpler simulation systems where mutation rate is encoded as a floating point value without a fine representation of the genotype-to-phenotype map and a realistic distribution of the effects of mutations. In a realistic setup, we expect mutations decreasing mutation rate to be scarce, and in particular rarer than mutations increasing it. Our system permits this realism, as fidelity systems are encoded on the genome and undergo mutations with an emergent, realistic distribution of effects, under the same rules than metabolic genes.

In a first attempt to disentangle the effects of decreased genetic drift and increased mutational supply in larger populations, we compared populations of different sizes at different time points to compensate for mutational supply. More specifically, populations 4 times smaller were evolved 4 times longer. We thus repeat the previous comparison, but analyze 4 million generations for populations of 256 individuals, 1 million generations for populations of 1024 individuals, and 200,000 generations for populations of 5041 individuals. Doing so, we found that the evolved mutation rates are mostly constant between population sizes for a given BMR value (Figure S4). Thus, we can not conclude that genetic drift is the sole or main driver: the fact that evolved mutation rates are lower when population size is increased may also be explained by an increased supply of gain-of-fidelity mutations. However, this correction is very conservative. An increase in population size also increases the fixation time of beneficial mutations. Moreover, as mutation rate evolves in this setup, it is hard to compare populations at equal opportunities for mutations. We are thus likely over-correcting, and this control can not be interpreted as evidence that genetic drift is not a limiting factor in mutation rate reduction.

We thus performed another control to assess the importance of both genetic drift and mutational supply: in populations which already evolved a low mutation rate, we strengthened genetic drift by switching to a smaller population size, and observed whether the mutation rate goes up. Starting from the genomes evolved in populations of 5041 individuals, we inoculated a clonal population of 256 individuals and let them evolve for an extra 4 million generations. We found that after the switch from 5041 to 256 individuals, the evolved mutation rates significantly goes up (Figure 4b). This suggests that genetic drift is an important factor, as mutational supply is less likely to be a limit in populations which *already* evolved alleles coding for a low mutation rate and are only challenged to *maintain* it. However, we also notice that mutation rates after the switch to a low population size remain lower than mutation rates of populations that always evolved at a low population size. This suggests that the supply of gain-of-fidelity mutations is also a limiting factor in mutation rate decrease.

### What is the impact of the explicit genomic representation on mutation rate evolution?

In previous experiments, mutation rate modifiers were encoded on the genome, following the same rules than genes directly contributing to fitness, and thus subject to a complex genotype-to-phenotype map. This permits a degree of realism absent in other models: individuals had to encode and maintain fidelity genes in their genome to lower their mutation rate, using genomic space that is limited. In addition, mutations reducing the mutation rate are rarer than mutations increasing it, and become even rarer the further the mutation rate decreased.

To explore the importance of theses modeling choices, relaxed these constraints by directly encoding mutation rates as floating point values without any genomic representation, as classically done in other models, but keeping the standard Aevol genotype-to-phenotype map for metabolic traits.

At each reproduction event, each individual has a fixed obability (1/500, independent of the current mutation ate value) of multiplicatively changing its mutation rate, upwards or downwards (with equal probabilities), by a constant factor (1.01). Genomes only encode metabolic genes (directly contributing to fitness), and the mutation rate defines the probability of a base-pair of this genome to mutate during reproduction. The BMR value is only used as the initial mutation rate value. We evolved populations of 1024 individuals for 1 million generations, and tested four different BMR values (10^-4^ , 10^-5^ , 10^-6^ and 10^-7^ ), with 10 replicates simulations for each value.

Populations with a mutation rate directly encoded as a floating point number evolved drastically lower mutation rates compared to when populations had to encode fidelity genes on the genome (Figure 5a). This is also true when looking at genomic mutation rate (Figure 5b). This shows that the genomic encoding of mutation rate matters, and that the evolved mutation rate value is not only determined by the selection pressure for mutational robustness.

**Fig. 5:**
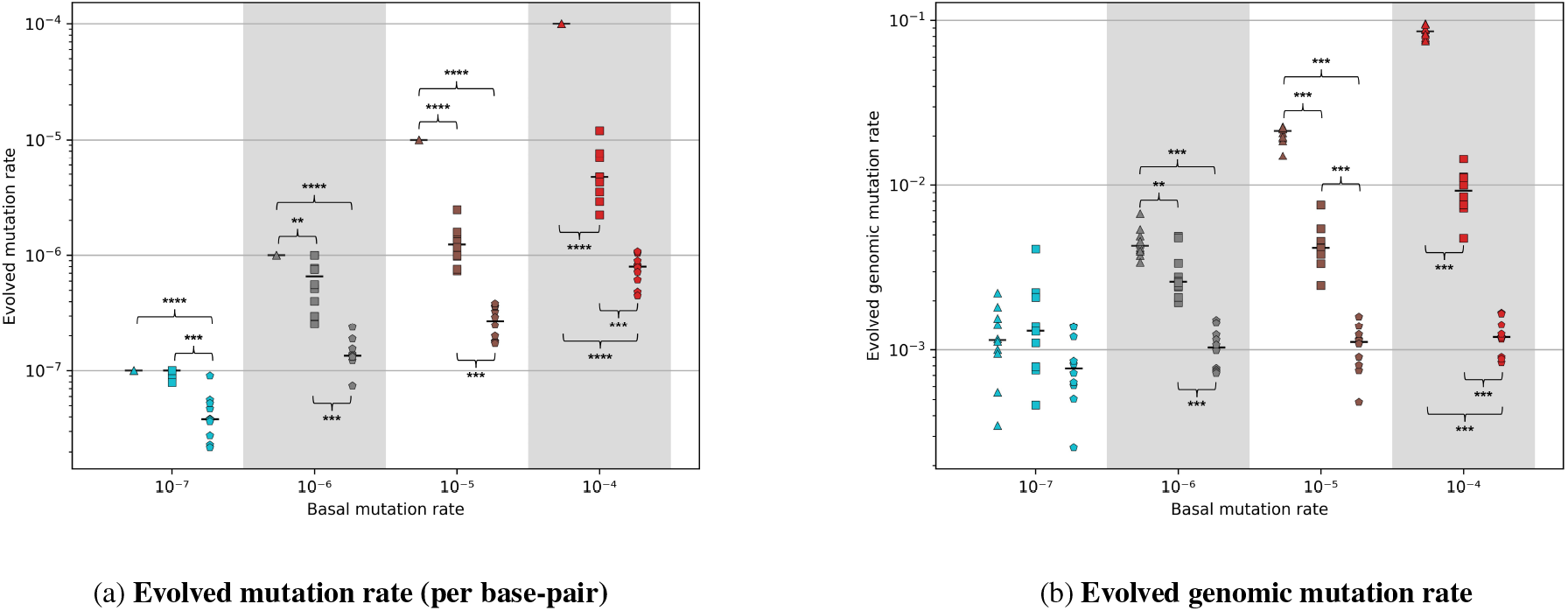
Evolution of the mutation rate represented as a floating point value. We compare populations which freely evolve a floating-point mutation rate by stepwise increments (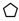) with the previously considered populations which can encode fidelity genes to decrease their mutation rate (□) or have a constant mutation rate (△). The population size is fixed at 1024 individuals, and each population evolved during 1 million generations. Four basal mutation rate (BMR) values are tested ranging from 10^-4^ to 10^-7^. For each BMR and each treatment, 10 replicates simulations are performed.

We also note that the populations with a floating point mutation rate did not evolve similar mutation rate between BMR conditions (Figure 5a), even though BMR values are only used as initial mutation rate values. However, they evolved similar genomic mutation rates (Figure 5b), in line with the postulate from Drake 1991. This confirms the idea that selection primarily acts to reduce the genomic mutation rate, which can be achieved by acting on both the mutation rate and the genome size. Depending on the initial mutation rate value, populations followed different evolutionary paths to reduce the genomic mutation rate to the same value.

Interestingly, we found that populations with a floating point mutation rate evolved a lower fitness than populations which can encode fidelity genes on the genome or control populations with a fixed mutation rate (Figure 6). This further confirms that selection pushes the mutation rate toward a suboptimal value, and shows that this sub-optimality is *even stronger* when the mutation rate reduction is not restricted by a realistic encoding of mutation rate limiting the accessibility of gain-of-fidelity mutations and inducing constraints on genome size.

**Fig. 6:**
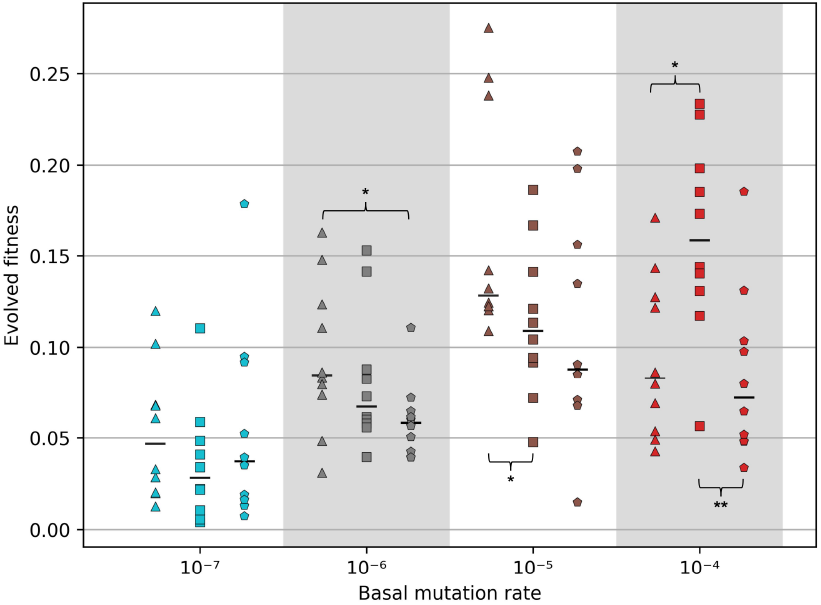
Evolved fitness when the mutation rate is represented as a floating point value. Populations have the ability to evolve their mutation rate by stepwise increments (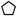), by encoding fidelity genes (□) or do not have it (△). The population size is fixed at 1024 individuals, and each simulation lasted 1 million generations. Four basal mutation rate (BMR) values are tested ranging from 10^-4^ to 10^-7^.

## Discussion

To summarize, we studied mutation rate evolution in a model with not only a complex fitness landscape, but also an explicit encoding of genes increasing replication fidelity following the same rules than metabolic genes directly impacting fitness. This permits several degrees of realism absent in most models, notably: (1) mutations affecting mutation rate have an emergent, realistic distribution of effects, with similar properties than mutations directly affecting fitness (epistasis, diminishing returns, more loss of functions than gain of functions, etc); (2) fidelity genes induce an indirect cost through increased genome size.

We observed that selection always pushes the mutation rate downwards, towards a value that is suboptimal for fitness trajectory. Population size is the main factor determining the amount of mutation rate reduction. We suggest that this link is not only explained by genetic drift, but also highlight the importance of access to gain-of-fidelity mutations. Finally, we recover a strong negative correlation between spontaneously evolved mutation rates and genome sizes.

### Selection favors sub-optimal mutation rate in spite of adaptation rate

Several theoretical models suggest that there exist an intermediate, optimal value for mutation rate, maximizing the rate of fitness increase for the populations (Kimura 1967; Levins 1967). This optimal would be the best compromise between maximizing the rate of adaptive mutations while minimizing the mutational load. This does not necessary imply that evolution of mutation rate will converge toward this value, but several models suggest that natural selection will indeed push mutation rate toward this optimal value (Leigh 1970, Bedau and Packard 2003). Yet, other works and our observations showed that selection is unable to do so and that evolution leads to suboptimal mutation rates (André and Godelle 2006; Clune et al. 2008; Good and Desai 2016), meaning that there exists another value that would have resulted in a higher fitness increase if mutation rate was constrained to this value.

This sub-optimality of spontaneously evolved mutation rate is thought to be linked to properties of the fitness landscape. The notion of fitness landscape refers to the mapping of the genotype to the fitness, and properties of this mapping (Wright 1932, Maynard Smith 1970, reviewed by Fragata et al. 2019). Empirical literature highlighted that ruggedness is a central feature of several experimentally-characterized fitness landscapes (Korona et al. 1994; Melnyk and Kassen 2011; Kouyos et al. 2012; Nahum et al. 2015). A fitness landscape is said to be rugged when composed of several fitness peaks connected by valleys. Epistasis, and in particular sign-epistasis, is an important source of ruggedness considered in the literature (Kouyos et al. 2012), although it does not systematically imply ruggedness (Weinreich et al. 2006).

The literature suggests that rugged fitness landscapes would drive evolution of mutation rate towards a lower than optimal value: this was shown by Clune et al. 2008 using simulations with digital organisms (Avida), and hypothesized in the discussion of Orr 2000. Briefly, the main argument is that on a rugged landscape, populations stay stuck on local fitness peaks for long periods of time between rare events of beneficial mutations which allow to reach a higher peak. During this time between two beneficial mutations, short-term selection against deleterious mutations pushes to a lower mutation rate. The more rugged the fitness landscape, the higher the waiting time between two beneficial mutations, and thus the lower the evolved mutation rate. Moreover, once the mutation rate starts decreasing, populations are less likely to generate beneficial mutations and will remain even longer on local fitness peaks (Weissman et al. 2009).

The genotype-to-phenotype map used in Aevol features strong sign-epistasis and a highly rugged fitness landscape: our findings are thus consistent with this literature. On a side note, the broad dispersion of fitness values between replicate populations (Figure 3) could precisely indicate that these populations are stuck on different local fitness peaks.

Another possible reason for sub-optimality of evolved mutation rates could be that fidelity genes act as selfish genetic elements, as they decrease their own probability of being removed by mutations, and therefore would tend to accumulate on the genome even when detrimental to the organism. This is close to the argument given by Lampert and Tlusty 2009, who argue that low mutation rate is a selfish trait. However, we saw that populations where mutation rate is directly encoded as a floating point value with fixed probability of change at each generation still evolved sub-optimal mutation rates, so this “ratchet” does not seem to be a necessary condition.

### Population size, genetic drift, and supply of gain-of-fidelity mutations

Within the context of a broad body of literature suggesting that selection mostly pushes mutation rate downwards, an hypothesis recently emerged regarding the role of genetic drift in this process. The drift-barrier hypothesis posits that genetic drift prevents the fixation of new gain-of-fidelity mutations beyond a threshold value determined by effective population size. This would explain a negative correlation between effective population size and mutation rate (Lynch et al. 2016).

Our simulation model permits to test such hypothesis. We do not directly control effective population size, but modified census population size and found that it strongly impacts mutation rate evolution (Figure 4).

However, population size does not only influence the strength of genetic drift, but also the total mutational supply for the population, and in particular the supply of gain-of-fidelity mutations. We highlight the role of this supply of gain-of-fidelity mutations, in particular through simulations where mutation rate is encoded as a floating point value, with a fixed distribution of mutations. Compared to these populations, it is increasingly difficult for populations encoding mutation rate on the genome to decrease their mutation rate because access to new gain-of-fidelity mutations become scarcer as fidelity systems improve. This is due to both diminishing-returns epistasis and a lower mutation rate.

These findings highlight the importance of considering a model which features not only a genotype-phenotype-fitness map with a realistic distribution of mutations and their effects, but also a “genotype to mutation rate” map, with also a realistic distribution of mutations affecting fidelity and their effects.

Sasaki and Nowak 2003 proposed the term *mutation landscape* to denote this equivalent of the genotype-phenotype-fitness mapping, but for mutation rate instead of traits directly impacting fitness.

Based on our modified version of Aevol, which features such mutation landscape in addition to the fitness landscape, we suggest that evolved mutation rate are not only determined by the strength of selection against deleterious mutations, but also by this mutation landscape.

### Genome size and the cost of fidelity

Whether a direct cost associated with replication fidelity plays a role in mutation rate evolution has been a long-standing question. While some empirical work suggest that the cost associated with replication fidelity observed in an RNA virus stems from decreased genetic diversity at the population scale (Vignuzzi et al. 2005; Pfeiffer and Kirkegaard 2005), other work suggest that fidelity genes can cause a direct fitness cost, for example due to time delay for DNA repair or extra energy consumption (Furió et al. 2005; Fitzsimmons et al. 2018). Theoretical literature suggests that direct phenotypic costs associated with fidelity systems could push the mutation rate towards an intermediate value (Ishii et al. 1989; André and Godelle 2006).

In this work, fidelity genes were not associated with an explicit direct fitness cost. But individuals had to allocate genomic space to encode fidelity genes, a space that could not be allocated to metabolic genes without also increasing the genome size. While individuals are not directly penalized for having a larger genome, previous results have shown that larger genomes are less robust and undergo more deleterious mutations, and therefore that high mutation rates select for shorter genomes (Eigen 1971; Knibbe et al. 2007).

This raises an interesting dilemma, as in our setup individuals need to encode more genes to decrease their mutation rate, but each extra base pair make them more prone to mutations. We found that even when the pressure on genome size was the strongest (in high BMR values, which correspond to the highest cost for genomic space), evolved populations still encoded fidelity genes – and a higher number than populations evolving with a lower BMR value. A possible explanation is that, after fidelity genes evolve, even through the genome size increased, the total number of mutations is lower than what would have happen without those fidelity genes and this extra genome size.

Our observations show that the cost for genomic space alone is not enough to drive the mutation rate toward an intermediate value.

## Model and Methods

### The Aevol model

Aevol is an individual-based, forward-in-time evolutionary simulations model. It is inspired by bacterial genomics, and have been used to study the evolution of the genome structure (Knibbe et al. 2007; Banse et al. 2024). Figure 7 pictures a coarse-grained representation of the model. The main features and parameters used in this study are detailed below. A more detailed explanation can be found in Aevol website (www.aevol.fr).

**Fig. 7:**
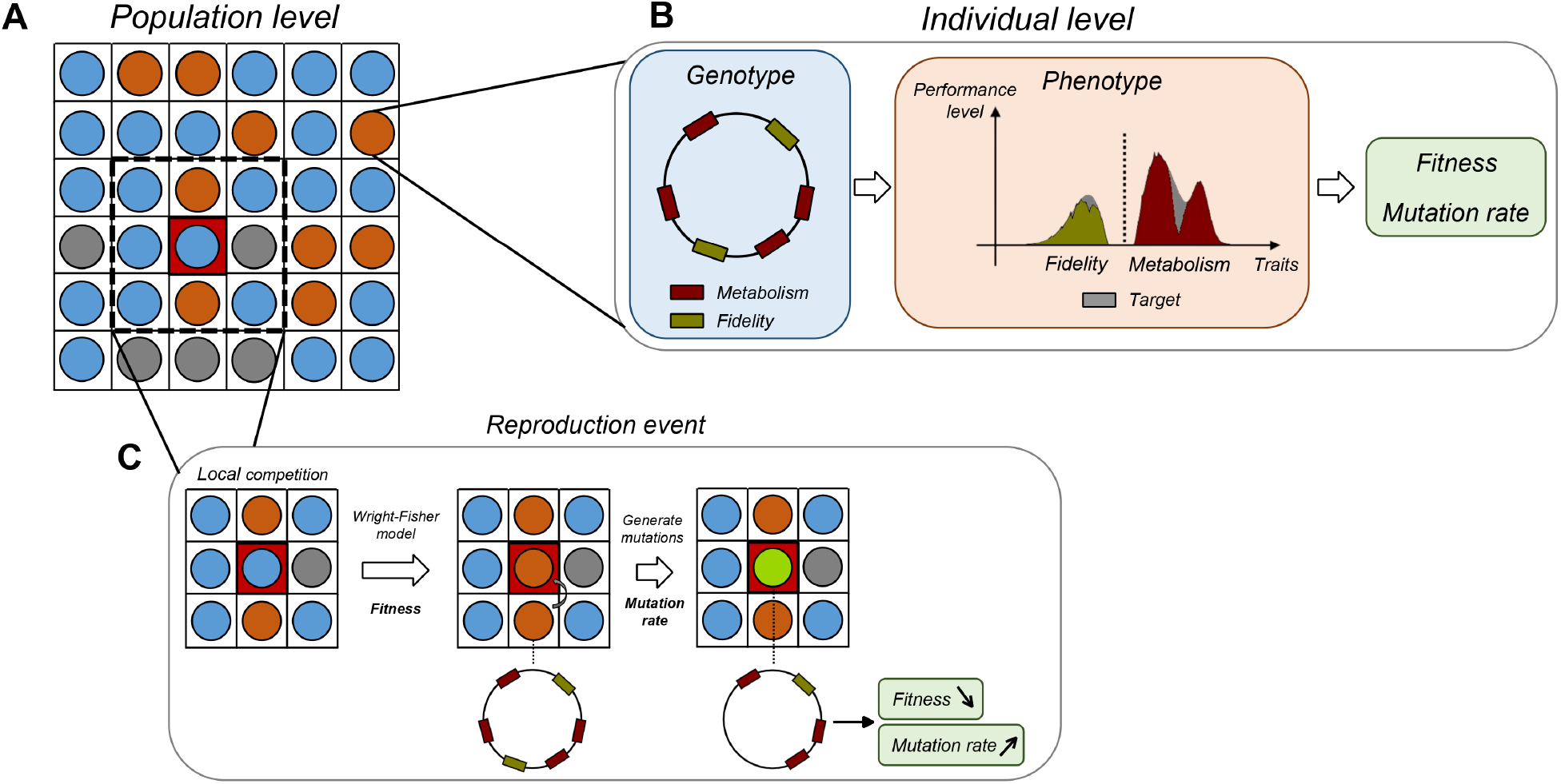
Schematic of the model (adaptation of Aevol with a variable mutation rate). A) A fixed number of individuals live on a 2D grid. B) Each grid cell contains an individual with a genome. Sequences of nucleotides codes for genes which can impact metabolism or fidelity. The phenotypes are compared with a predefined target (in gray, which correspond to perfect metabolic and fidelity phenotypes) to determine fitness and mutation rate of the individual. C) At each generation, the population is replaced in two steps. First, following a Wright-Fisher model, each individual is replaced by an offspring of another individual chosen locally in the 3 × 3 neighborhood based on fitness. Second, the offspring may undergo mutations whose probability depends on the mutation rate of the parent. In the example displayed, the offspring undergoes a large deletion, resulting in the loss of one metabolic gene and one fidelity genes, thereby decreasing its fitness and increasing its mutation rate.

#### Demographics

The model features a fixed population size with spatial structure (two-dimensional toroidal lattice), and Wright-Fisher reproduction scheme, with local reproduction: at each simulation step, each individual is replaced by the offspring of an individual in the 3 × 3 neighborhood. The probability of reproduction within the neighborhood is proportional to fitness, computed as explained further below.

#### Genotype-to-phenotype

Each individual has a circular genome, composed of 0 and 1, from which a phenotype can be computed. The first step, which represents the transcription, consists in searching for two types of sequence motifs on the genome: the promoters, defined by similarity to a consensus sequence, which mark the beginning of a transcript, and the terminators, defined as inverted repeats which form a hairpin structure and mark the end of the transcript. The second step, which represents the translation, consists in finding, inside the previously identified transcripts, a Shine-Dalgarno-like sequence followed by a START codon, which marks the beginning of a gene. Then, the sequence is read 3 bases at a time and interpreted as codons following a predefined genetic code, until a STOP codon is found in frame with the START codon. The sequence of codons is converted to a protein, and the set of all proteins defines a phenotype, following an abstract mathematical encoding described in details by Rutten et al. 2019. Briefly, a phenotype is represented as a [0, 1] → ℝ function: the x-axis corresponds to the functional space (continuous set of all possible biological traits) and the y-axis corresponds to the realization level (how much a given trait is performed by the considered genome).

#### Phenotype-to-fitness

The fitness of an individual depends on the distance between its phenotype and a target function, which represents the ideal phenotype in the specified environment. In the present study, the target function is defined by the sum of three gaussian distributions whose parameters are given in Table S1.

The fitness is calculated as:

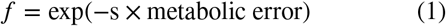

where s is the selection pressure parameter, and the metabolic error is the distance between the phenotype of the individual and the target function.

#### Mutations

At each reproduction event, the individual can undergo local mutations or chromosomal rearrangements, which may affect the phenotype, and ultimately can change the fitness or mutation rate. The phenotype of a mutant offspring is computed from its genotype following the rules described above. The distribution of the effects of mutations is thus an emergent property of the model, and not an input parameter.

The mutation rate is defined per base-pair. Different types of mutations (small indels, point mutation, large duplication, etc) can have different mutation rate values, although we choose to keep the same value for all mutation types in this study. The size distribution of small indels is uniform from 1 to 6 base-pairs. The size distribution of chromosomal rearrangements is uniform from 1 to the total genome size.

### Specific adaptations of the Aevol model for this study

#### Incorporation of a variable mutation rate

In the basic version of Aevol, the entire axis of traits (from 0 to 1) represents classical traits which directly impact the phenotype and are thus under first order selection (*metabolism*). In our modified version, the trait axis is divided in two parts: one part contributing to the fidelity of replication (from 0 to 0.5), and the other part to the metabolism (from 0.5 to 1). The former part does not impact fitness computation, but impacts mutation rate.

#### Mutation rate computation

In this setup where populations can encode fidelity genes to decrease their mutation rate, the mutation rate is calculated as:

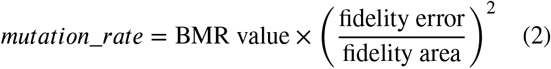

Where the fidelity error is the distance, from 0 to 0.5 on the functional space axis, between the phenotype of the individual and the target function, and the fidelity area is the area of the target function from 0 to 0.5.

In the control where populations have a fixed mutation rate, this mutation rate directly equals the BMR value.

### Experimental setups

#### Initial conditions for evolutionary simulations

Unless specified otherwise, all simulations were initialized with a clonal population of individuals with a single metabolic gene, randomly obtained. There are two reasons for starting with a genome containing a metabolic gene rather than a fully random sequence : i) the apparition of the first metabolic gene can take some time during which nothing evolutionary interesting happens; and ii) under high BMR values, when a genome does not start with at least one metabolic gene, we noticed that mutational bias often reduces it to 1 base-pair (the hard limit), after which the individuals stay stuck with empty genomes.

#### Evolution of replication fidelity (Figures 1, 2, 3)

In this setup, populations either have or do not have the ability to reduce their mutation rate. The population size is fixed at 1024 individuals and the simulations lasted 4 million generations. Seven BMR values were tested (10^-4^ , 5.10^-5^ , 10^-5^ , 5.10^-6^ , 10^-6^ , 5.10^-7^ and 10^-7^ ), with ten replicate simulations for each BMR value. Each replicate has the same initial conditions except for a random number seed, which impact the outcome of the stochastic events. In total, 140 simulations were performed.

#### Evolution with a lower bound for mutation rate (Figure S3)

This setup is similar to “Evolution of replication fidelity”, but with a lower bound for the mutation rate. The mutation rate is equal to the lower bound if the computed mutation rate is below it. Two BMR values were tested (10^-4^ and 5.10^-5^ ), with ten replicate simulations for each BMR value. The lower bound of the BMR value 10^-4^ is set to 10^-5^ , and the BMR value 5.10^-5^ to 5.10 ^-6^. In total, 20 simulations were performed.

#### Effect of population size (Figure 4a)

This setup is similar to “Evolution of replication fidelity”, but two other population sizes are tested, 256 or 5041 individuals, which correspond to a grid size of 16 by 16 or 71 by 71. All populations have the ability to reduce their mutation rate. Only four BMR values were tested (10^-4^ , 10^-5^ , 10^-6^ and 10^-7^ ) because of the computation load, with ten replicate simulations for each BMR value. In total, 80 simulations were performed.

#### Switching population size during the simulation (Figure 4b)

In this setup, we extracted the genome of the individual with the highest fitness at the end of each replicate simulation with 5041 individuals in the setup “Effect of population size”. We used each genome to seed a clonal population of 256 individuals for an additional phase of evolution of 4 million generations. Three BMR values were tested (10^-4^ , 10^-5^ and 10^-6^ ). In total, 30 simulations were performed.

#### Encoding mutation rate as a floating point value (Figures 5a, 5b, 6)

In this setup, populations have the ability to change their mutation rate with a fixed probability (1/500), upwards or downwards (with equal probabilities), by a constant factor (1.01). The population size is fixed at 1024 individuals and the simulations lasted 1 million generations. Four BMR values were tested (10^-4^ , 10^-5^ , 10^-6^ and 10^-7^ ), with ten replicate simulations for each BMR value. In total, 40 simulations were performed.

### Data display

After the simulations ended, we reconstructed the ancestral lineage of the individual with the highest fitness at the end. On this lineage, we extracted all the associated features (fitness, mutation rate etc.) at each generation. In almost all figures, the data points correspond to the mean value of the feature during the last 100,000 generations of the lineage. The only exception being in Figure 4b for the mutation rate at the start which is taken at generation 0 only.

## Code and data availability

We based our code on commit 87543ed9, branch aevol-9, in the public repository of Aevol, which is a free software distributed under the GPL. The modified version of the simulator, as well as the raw simulation data, the code used to analyze and display these data, and the processed data, are available on Zenodo: 10.5281/zenodo.20931661.

## Acknowledgments

We express our gratitude to Dule Misevic, François Taddei and Guillaume Beslon for inspiring discussions around mutation rate evolution, to David Parsons and Guillaume Beslon for discussions related to the Aevol system, and to the High Performance Computing center from the University of Grenoble (GRICAD) for providing the computational power needed for this study.

## Funding

This work received financial support from MIAI@Grenoble Alpes (ANR-19-P3IA-0003).

## Supplementary materials

**Fig. S1:**
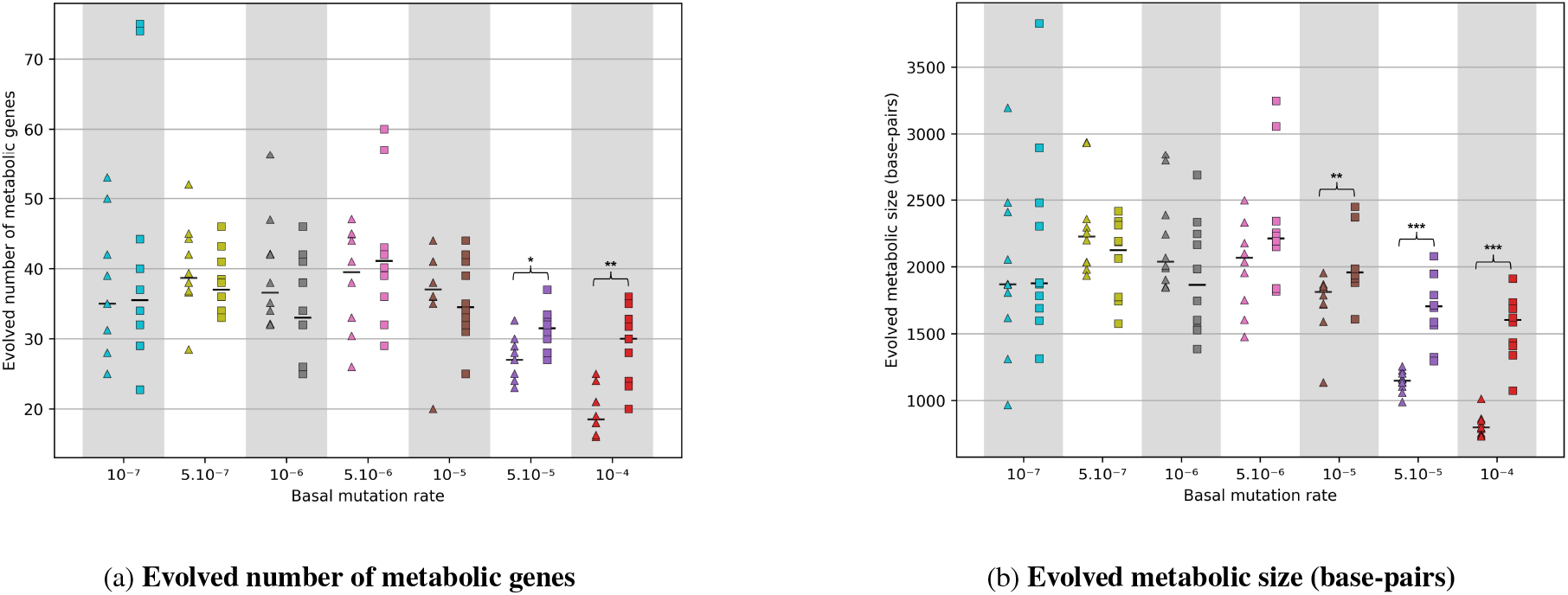
Evolved metabolism. Populations either have the ability to reduce their mutation rate (□) or do not have it (△). Seven basal mutation rate (BMR) values are tested ranging from 10^-4^ to 10^-7^. For each BMR and each treatment, 10 replicates simulations are performed. Each simulation lasted 4 million generations. Each data point corresponds to the mean for a replicate simulation during the last 100,000 generations (see data display section in Model and Methods for more details), and the horizontal bar represents the median among replicate simulations.

**Fig. S2:**
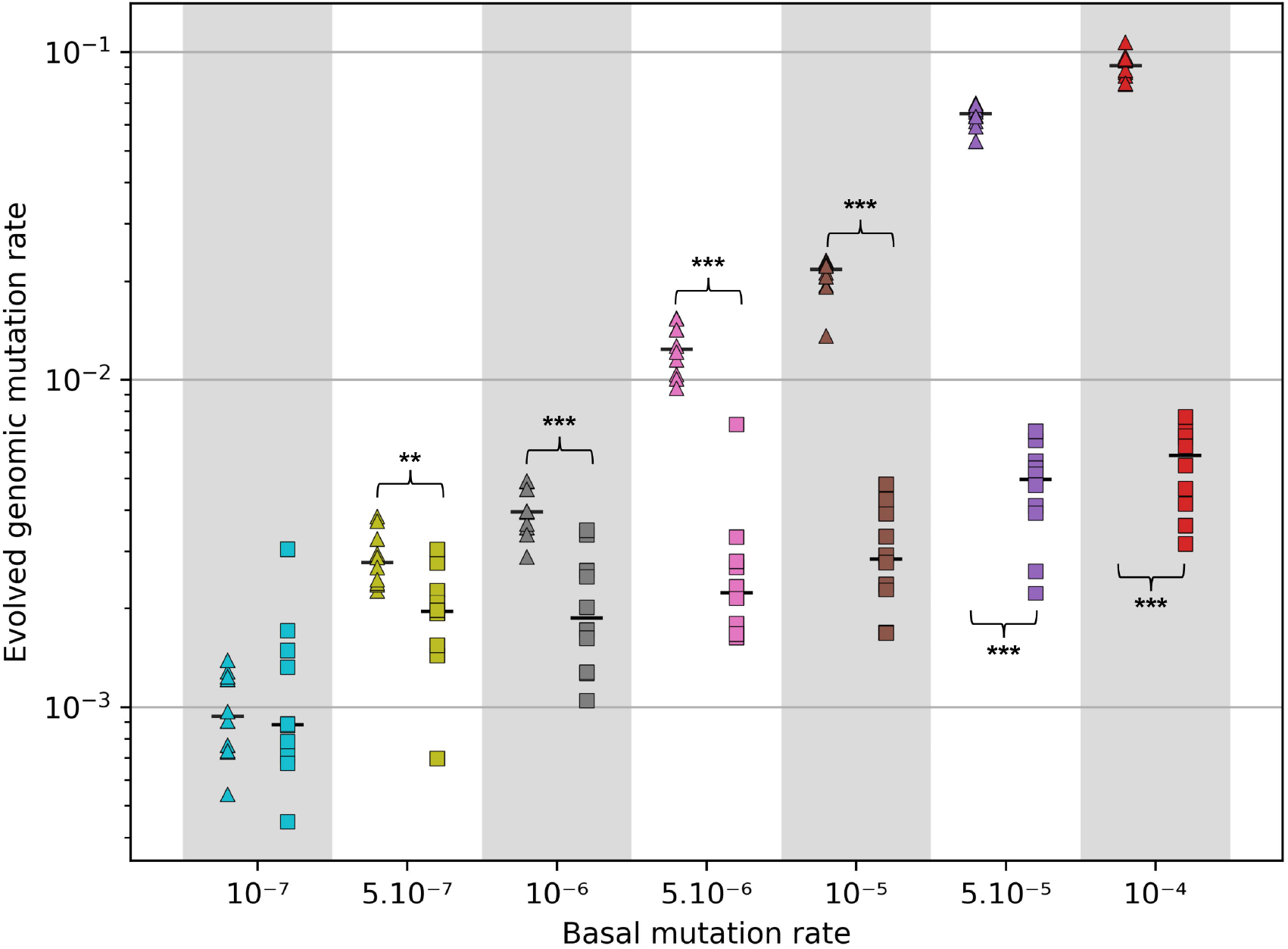
Evolved genomic mutation rate. Populations either have the ability to reduce their mutation rate (□) or do not have it (△). Seven basal mutation rate (BMR) values are tested ranging from 10^-4^ to 10^-7^. For each BMR and each treatment, 10 replicates simulations are performed. Each simulation lasted 4 million generations. Each data point corresponds to the mean for a replicate simulation during the last 100,000 generations (see data display section in Model and Methods for more details), and the horizontal bar represents the median among replicate simulations.

**Fig. S3:**
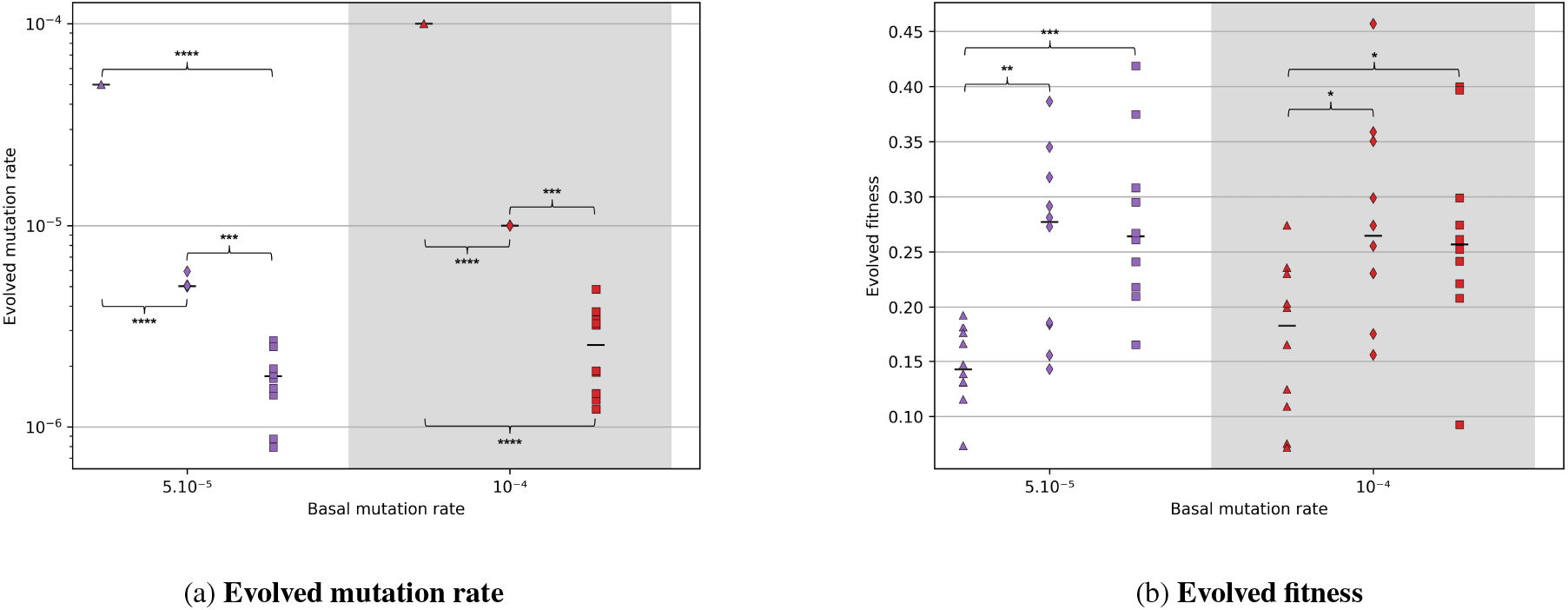
Evolved mutation rate and fitness with a lower-bounded mutation rate. Populations either have the ability to reduce their mutation rate (□), have a lower-bounded mutation rate of one order of magnitude below their BMR value (⋄) or don’t have the ability to reduce the mutation rate (△). Two basal mutation rate (BMR) values are tested (10^-4^ and 5.10^-5^). For each BMR and each treatment, 10 replicates simulations are performed. Each simulation lasted 4 million generations. Each data point corresponds to the mean for a replicate simulation during the last 100,000 generations (see data display section in Model and Methods for more details), and the horizontal bar represents the median among replicate simulations.

**Fig. S4:**
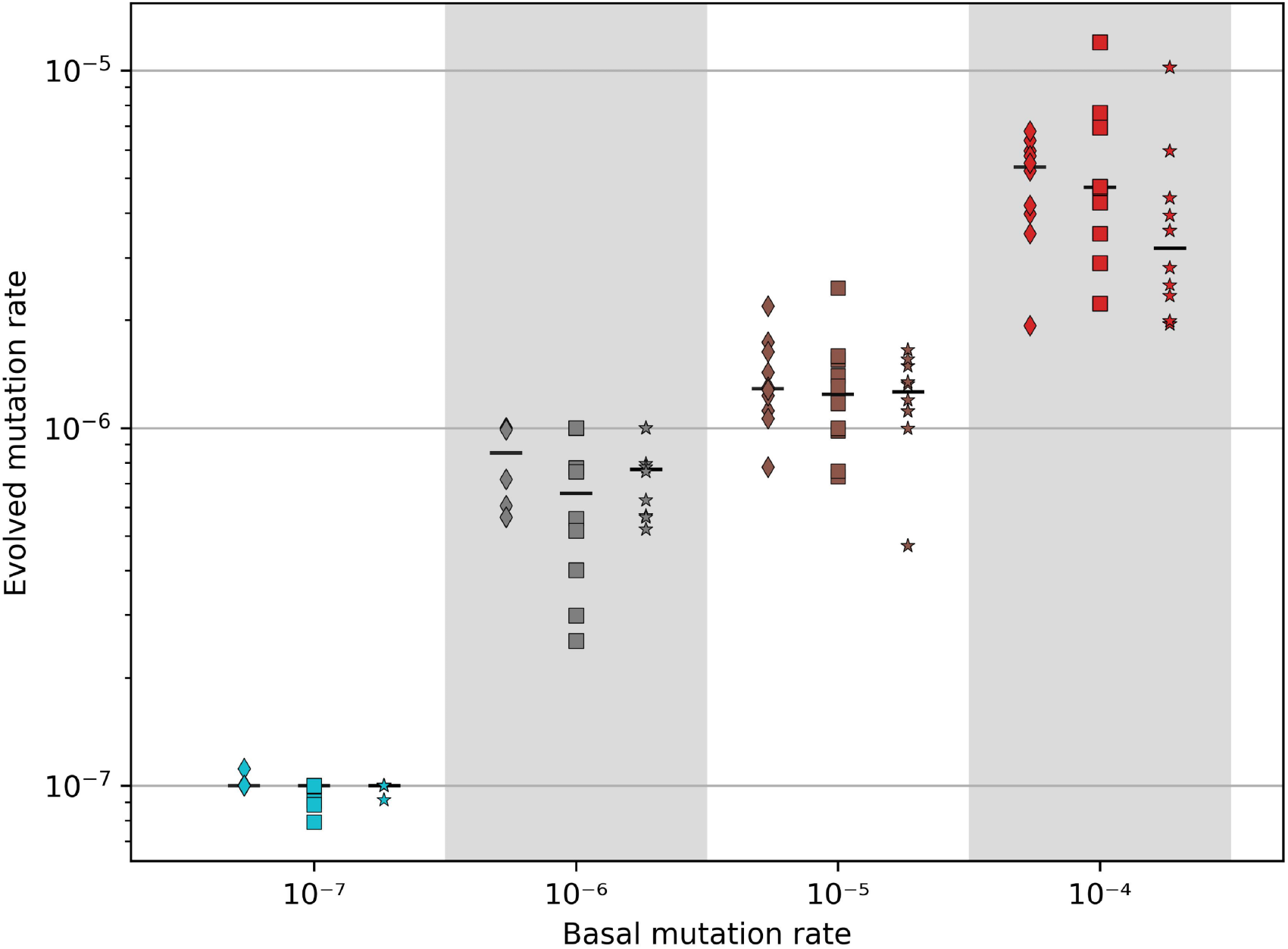
Evolved mutation rate, comparing populations with the same population size × nb generations product. All populations have the ability to reduce their mutation rate. Populations of 256 individuals (⋄) evolved for 4 million generations, populations of 1024 individuals (□) evolved for 1 million generations and populations of 5041 individuals (⋆) evolved for 200,000 generations. Four basal mutation rate (BMR) values are tested ranging from 10^-4^ to 10^-7^. For each BMR and each treatment, 10 replicates simulations are performed. Each data point corresponds to the mean for a replicate simulation during the last 100,000 generations (see data display section in Model and Methods for more details), and the horizontal bar represents the median among replicate simulations.

**Table S1:** Aevol parameters. SEED_VALUE, GRID_WIDTH, BMR_VALUE are dependent on the experimental setups.

| Section | Parameter | Value |
| --- | --- | --- |
| **1. Initial setup** |  |  |
|  | SEED | <SEED_VALUE> |
|  | GRID_SIZE | <GRID_WIDTH> <GRID_WIDTH> |
|  | CHROMOSOME_INITIAL_LENGTH | 200 |
| **2. Selection** |  |  |
|  | SELECTION_SCHEME | fitness_proportionate 1000 |
| **3. Mutation rates** |  |  |
|  | POINT_MUTATION_RATE | <BMR_VALUE> |
|  | SMALL_INSERTION_RATE | <BMR_VALUE> |
|  | SMALL_DELETION_RATE | <BMR_VALUE> |
|  | MAX_INDEL_SIZE | 6 |
| **4. Rearrangement rates (w/o alignments)** |  |  |
|  | DUPLICATION_RATE | <BMR_VALUE> |
|  | DELETION_RATE | <BMR_VALUE> |
|  | TRANSLOCATION_RATE | <BMR_VALUE> |
|  | INVERSION_RATE | <BMR_VALUE> |
| **6. Target function** |  |  |
|  | ENV_SAMPLING | 300 |
|  | ENV_ADD_GAUSSIAN | 1.2 0.52 0.12 |
|  | ENV_ADD_GAUSSIAN | -1.4 0.5 0.07 |
|  | ENV_ADD_GAUSSIAN | 0.3 0.8 0.03 |
|  | MAX_TRIANGLE_WIDTH | 0.033333333 |
| **7. Recording** |  |  |
|  | CHECKPOINT_STEP | 50000 |
|  | RECORD_TREE | ON 50000 |
|  | STATS_BEST | ON 1000 |
|  | STATS_POP | OFF 1000 |
| **8. Environment variation** |  |  |
|  | ENV_VARIATION | none |

## Notes

### Competing Interest Statement

The authors have declared no competing interest.

